# Metformin induces lipogenic differentiation in myofibroblasts to reverse mouse and human lung fibrosis

**DOI:** 10.1101/401265

**Authors:** Vahid Kheirollahi, Roxana M. Wasnick, Valentina Biasin, Ana Ivonne Vazquez-Armendariz, Xuran Chu, Alena Moiseenko, Astrid Weiss, Jochen Wilhelm, Jin-San Zhang, Grazyna Kwapiszewska, Susanne Herold, Ralph T. Schermuly, Werner Seeger, Andreas Günther, Saverio Bellusci, Elie El Agha

## Abstract

Idiopathic pulmonary fibrosis is a fatal, incurable lung disease in which the intricate alveolar network of the human lung is progressively replaced by fibrotic scars, eventually leading to respiratory failure. Myofibroblasts are the effector cells that lead to abnormal deposition of extracellular matrix proteins and therefore mediate fibrotic disease not only in the lung but also in other organs. Emerging literature suggests a correlation between fibrosis and metabolic alterations in IPF. In this study, we show that the first-line antidiabetic drug, metformin, exerts potent antifibrotic effects in the lung by modulating metabolic pathways, inhibiting TGFβ1 action, suppressing collagen formation, activating PPARγ signaling and inducing lipogenic differentiation in lung myofibroblasts derived from human patients. Using genetic lineage tracing in a murine model of lung fibrosis, we show that metformin alters the fate of myofibroblasts and accelerates fibrosis resolution by inducing myofibroblast-to-lipofibroblast transdifferentiation. Detailed pathway analysis showed that the reduction of collagen synthesis was largely AMPK-dependent, whereas the transdifferentiation of myo- to lipofibroblasts occurred in a BMP2-PPARγ-dependent fashion and was largely AMPK-independent. Our data report an unprecedented role for metformin in lung fibrosis, thus warranting further therapeutic evaluation.

## Introduction

Idiopathic pulmonary fibrosis (IPF) is a fatal lung disease of unknown etiology. This disease is more common among the elderly and the average survival rate following diagnosis is only 2-3 years^1^. Histopathological examination of IPF lungs typically reveals extensive alveolar scarring; i.e. replacement of normal alveoli by fibrous scars containing myofibroblasts. The latter cells are considered to be the main source of excessive extracellular matrix (ECM) protein deposition, particularly collagen^2^, not only in IPF lungs but also in fibrosis of other organs. Due to its progressive nature and since the process of scar formation is part of natural wound healing, IPF is widely regarded as an aberrant wound healing response to repetitive epithelial injury^3^.

The cellular source of myofibroblasts has been a subject of debate in recent years. The paradigm is that identifying the precursor cell of the myofibroblast might pave the way for preventive and/or therapeutic, selective intervention in IPF patients. In this context, it is suggested that the myofibroblast pool is heterogeneous, and derives from multiple sources such as resident fibroblasts, circulating fibrocytes, perivascular mesenchymal cells and alveolar epithelial cells^4–7^. Using genetic lineage tracing, we have recently identified the resident lipid-droplet-containing interstitial fibroblast, or lipofibroblast, as a precursor cell for the myofibroblast in the bleomycin model of lung fibrosis in mice^8^. We also found that myofibroblasts retain their plasticity and are able to revert to the type 2 alveolar epithelial cell-supportive lipofibroblast fate during recovery^8^. We finally showed that peroxisome proliferator-activated receptor gamma (PPARγ) agonist, rosiglitazone, induces lipogenic differentiation and inhibits transforming growth factor beta 1 (TGFβ1)-induced myogenic differentiation in primary human lung fibroblasts^8^.

The therapeutic effect of rosiglitazone in the murine model of bleomycin-induced pulmonary fibrosis is well established^9,10^. Rosiglitazone belongs to the thiazolidinedione class of PPARγ agonists and is a potent antidiabetic agent. Ongoing research regarding its role in suppressing inflammation^11^, inducing apoptosis and suppressing tumor growth^12^ in parallel to inhibiting TGFβ1 signaling^8^, makes it a good therapeutic candidate for IPF patients. However, increased risk of ischemic heart disease reported in some clinical trials reduces the enthusiasm for implementing it in the management of IPF^13^.

Interestingly, IPF is associated with metabolic disorders. For example, a recent report has shown that IPF lungs display alterations in several metabolites linked to energy consumption^14^. In addition, there is evidence suggesting that several types of lipid molecules present in blood plasma could be used as biomarkers for IPF^15^. One study has even suggested that type 2 diabetes might be a risk factor for developing IPF^16^. Although it remains unclear whether these metabolic abnormalities represent a cause or consequence of the disease, the link between fibrosis and metabolic alterations in the lung raises the question whether antidiabetic drugs can be good candidates for antifibrotic therapy. Metformin is another antidiabetic drug that has been extensively used to manage diabetic patients. It inhibits gluconeogenesis in the liver and increases peripheral glucose utilization by sensitizing cells to insulin. In fact, metformin and rosiglitazone have been used in combination to treat patients with type 2 diabetes. Both agents decrease the amount of glucose produced by the liver and absorbed by the intestine.

Many studies have reported the therapeutic effects of metformin in non-diabetic diseases such as non-small-cell lung cancer^17^, prostate cancer^18^ and cardiovascular diseases^19^. Moreover, it has been suggested that intraperitoneal administration of metformin attenuates bleomycin-induced lung fibrosis in mice via NADPH oxidase 4 (NOX4) suppression^20^. A more recent report has shown that metformin accelerates resolution of bleomycin-induced pulmonary fibrosis, suggesting activation of AMP-activated protein kinase (AMPK) as key underlying signaling event, leading to downregulation of alpha smooth muscle actin (*ACTA2*) and collagen, and increasing myofibroblast autophagy and ECM turnover^21^.

In the current study, we hypothesized that metformin accelerates fibrosis resolution by inducing lipogenic differentiation in lung fibroblasts, while inhibiting TGFβ1-mediated myogenic differentiation. To test this hypothesis, we used primary cultures of human IPF-derived lung fibroblasts, cultures of precision-cut lung slices (PCLS) derived from human IPF patients, and genetic lineage tracing in the context of the bleomycin model of lung fibrosis in mice. We show that metformin accelerates fibrosis reversal by altering the fate of myofibroblasts in the lung. Mechanistically, metformin induces lipogenic differentiation in myofibroblasts via a mechanism involving bone morphogenetic protein 2 (*BMP2*) upregulation and PPARγ activation, and inhibits TGFβ1-induced collagen production via AMPK activation. Our data highlight the potential for using metformin to treat IPF patients.

## Methods

### Human-derived specimens

Human lung tissue samples and primary lung fibroblasts were obtained through the European IPF registry (eurIPFreg) at the Universities of Giessen and Marburg Lung Center, member of the German Center for Lung Research^22^. Written consent was obtained from each patient and the study was approved by the ethics committee of Justus-Liebig University Giessen.

### Animal experiments

All animals were housed under specific pathogen-free (SPF) conditions with free access to food and water. At 11-12 weeks of age, *Acta2-Cre-ERT2; tdTomato^flox^* males were subjected to intratracheal instillation of saline or bleomycin (0.8 U/kg body weight) (Sigma-Aldrich) using a micro-sprayer (Penn-Century, Inc.) at the Ludwig Boltzmann Institute in Graz, Austria. Five days after bleomycin instillation, mice were fed tamoxifen-containing chow (400 mg/kg food, Envigo) for nine days. Metformin (1.5 mg/mL) or vehicle (PBS) was supplied via drinking water at day 14 after bleomycin instillation. Lungs were harvested on day 28. All animal experiments were approved by the local authorities (Austrian Ministry of Education, Science and Culture; BMWFW-66.010/0043-WF/V/3b/2016) and performed in accordance with the EU directive 2010/63/EU.

### Cell culture

Primary lung fibroblasts derived from 12 IPF patients were maintained in Dulbecco’s Modified Eagle’s Medium (DMEM) (Invitrogen) supplemented with 10% bovine calf serum (BCS, Gibco) at 37 °C and 5% CO_2_. Cells between passages 3 and 7 were used for the experiments. Briefly, 3x10^5^ cells were seeded per well in 6-well plates (Greiner Bio-One). The next day, cells were starved for 24 h and then treated with different compounds. For imaging experiments, cells were cultured in 4-well chamber slides (Sarstedt) at a density of 75,000 cells per well. Cells were treated with metformin (Ratiopharm), pirfenidone (Cayman Chemical Company), nintedanib, selumetinib or GSK621 (all from Selleckchem), recombinant human TGFβ1 (rhTGFβ1) or rhBMP2 (both from R&D Systems). Table 1 summarizes treatment conditions. Control cells were treated with the corresponding solvents as recommended by the manufacturer (same volumes as treated groups).

**Table 1.** Treatment conditions of primary human lung fibroblasts.

| Compound | Final concentration | Solvent |
| --- | --- | --- |
| GSK621 | 10 μM* | DMSO |
| Metformin | 1 mM, 5 mM, 10 mM | DMEM |
| Nintedanib | 1 μM* | DMSO |
| Pirfenidone | 0.3 mg/mL* | Ethanol |
| Selumetinib | 5 μM* | DMSO |
| rhTGFβ1 | 2 ng/mL | rhTGFβ1 reconstitution buffer |
| rhBMP2 | 50 ng/mL* | rhBMP2 reconstitution buffer |

\* Dosage was chosen based on a literature search and the dose showing highest efficacy was selected.

### Precision-cut lung slices

Fresh lung specimens were obtained from 4 IPF patients that underwent lung transplantation. Precision-cut lung slices were prepared in two ways: lung specimens were cut into strips (length: 2-3 cm, thickness: 3-5 mm) and later chopped into 200 µm-thick slices using a McIlwain Tissue Chopper (Campden Instruments Ltd.); or lung tissues were gently injected with 1.5% low-melting agarose (Roth) and cut using a vibratome (Thermo Fisher Scientific) into 400 µm-thick slices. Five to six PCLS were cultured in 5 mL of DMEM supplemented with 10% BCS at 37 °C and 5% CO_2_ for five days. Cultures were treated with different agents at the beginning of the culture process.

### siRNA transfection

siGENOME PRKAA1 siRNA (D-005027-01-0002) and the corresponding scrambled siRNA (siGENOME Non-Targeting siRNA #2 (D-001210-02-05)) were obtained from Dharmacon. When cells reached 50-60% confluence in 6-well plates, they were transfected using lipofectamine RNAiMAX (Invitrogen) according to manufacturer’s instructions (25 pmol per well, 7.5 µL lipofectamine per well). The culture medium was replaced 24 h after transfection. Seventy-two hours after transfection, the culture medium was replaced by either fresh medium or fresh medium supplemented with 5 mM metformin.

### RNA extraction and quantitative real-time PCR

Total RNA extraction was performed using RNeasy mini or micro kits (Qiagen) and cDNA synthesis was carried out using Quantitect reverse transcription kit (Qiagen) according to manufacturer’s instructions. Quantitative real-time PCR (qPCR) was performed using PowerUp SYBR green master mix (Applied Biosystems) and LightCycler 480 II machine (Roche Applied Science). Porphobilinogen deaminase (*PBGD*) was used as a reference gene. Data are presented as expression levels relative to *PBGD* using the 2^-ΔCT^ method. Primer sequences are shown in Table 2.

**Table 2.**
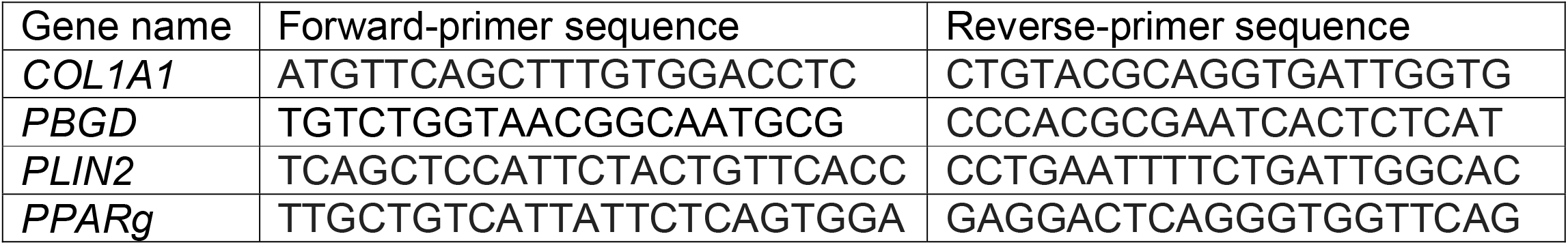
Primer sequences for qPCR.

### Protein extraction and Western blotting

Total protein lysates were prepared using RIPA buffer (Santa Cruz) according to manufacturer’s instructions. Proteins were separated by sodium dodecyl sulfatepolyacrylamide gel electrophoresis (SDS-PAGE) followed by blotting on polyvinylidene fluoride (PVDF) membranes (Thermo Fisher Scientific) using Trans-Blot SD semi-dry transfer cell (Bio-Rad Laboratories). Antibodies against AMPK (Abcam, 1:2500), PPARγ (H-100) (Santa Cruz, 1:1000), phospho-PPARγ (Merck Millipore, 1:500), p44/42 mitogen-activated protein kinase (MAPK, a.k.a. extracellular signal-regulated kinases 1 and 2, ERK1/2) (Cell Signaling, 1:1000) and phosphop44/42 MAPK [(ERK1) (Tyr204)/(ERK2) (Tyr187) (D1H6G)] (Cell Signaling, 1:1000) were used overnight at 4°C. Beta-actin (ACTB) was used as a loading control (Biolegend, 1:2500). HRP-conjugated anti-rabbit IgG (Promega, 1:5000) and HRPconjugated anti-mouse IgG (H+L) (Promega, 1:5000) were used as secondary antibodies for 1 h at room temperature (RT). Subsequently, membranes were covered with AceGlow chemiluminescence substrate (Peqlab) and imaged immediately using ChemiDoc XRS+ (Bio-Rad Laboratories).

### Staining for lipid-droplet accumulation

For time-lapse imaging, live cells were treated with different agents and HCS LipidTOX red neutral lipid dye (Invitrogen, 1:200) was immediately added. Cells were then placed in the incubation chamber (37 °C and 5% CO_2_) of DMI6000 B live imaging microscope (Leica). Images were acquired every 1 h. In other cases, cells were fixed using 2% paraformaldehyde (PFA, Roth) for 20 min followed by washing with PBS (Gibco). A mixture of diluted LipidTOX (1:200) and Hoechst (1:5000) in PBS was then used to stain fixed cells. Subsequently, slides were mounted using ProLong Gold Antifade Reagent (Molecular Probes).

### Hematoxylin and Eosin staining

Human-derived PCLS or mouse lung tissues were fixed using 4% PFA followed by embedding in paraffin. Paraffin blocks were sectioned into 5 µm-thick slices and placed on glass slides. Following deparaffinization, lung sections were stained with hematoxylin (Roth) for 2 min, washed with running tap water for 10 min and then stained with eosin (Thermo Fisher Scientific) for 2 min.

### Masson’s trichrome staining and fibrosis quantification

Collagen staining was performed using Masson’s trichrome staining kit according to the protocol recommended by the manufacturer (Thermo Fisher Scientific). Fibrosis was assessed by semi-automated quantification using VIS Image Analysis Software (Visiopharm). In brief, the algorithm calculates the percentage of the area covered by collagen fibers relative to the area covered by lung tissue (excluding airways and airspaces).

### Total collagen assay

In order to assess total collagen levels, 10 mg of each tissue sample was subjected to the total collagen assay kit according to manufacturer’s instructions (Biovision).

### Immunofluorescence

Following deparaffinization, slides underwent antigen retrieval using citrate buffer (Vector Laboratories) for 15 min followed by cooling on ice for 20 min. Slides were then blocked with 3% bovine serum albumin (BSA, Jackson Immunoresearch Laboratories) in PBS for 1 h at RT. Anti-collagen 1 A1 (anti-COL1A1) antibodies (Rockland, 1:200) and goat anti-rabbit antibodies (Life Technologies, 1:500) were used for immunofluorescence. Slides were finally mounted with ProLong Gold Antifade Reagent containing DAPI (Molecular Probes).

Human PCLS were fixed in 4% PFA for 2 h and stored in PBS containing 0.02% sodium azide. Fixed PCLS were blocked with 3% BSA in PBS for 2 h at RT, stained with anti-COL1A1 antibodies (Rockland, 1:200) overnight at 4°C and washed with PBS (three times, 30 min each). A mixture of Alexa Fluor 488-conjugated anti-rabbit antibodies (Invitrogen, 1:200), HCS LipidTOX red neutral lipid dye (Invitrogen, 1:200) and Hoechst (1:5000) was added for 3 h at RT. After washing, PCLS were placed in glass-bottomed 4-well micro slides (Ibidi) containing PBS and imaged by confocal microscopy (Leica TCS SP5).

For ACTA2 immunostaining following PFA fixation, samples were blocked with 3% BSA in PBS supplemented with 0.4% Triton-X (Sigma-Aldrich) for 1 h at RT and then stained with FITC-conjugated mouse monoclonal anti-ACTA2 antibodies (Sigma-Aldrich,1:200) overnight at 4°C. Slides were finally mounted with ProLong Gold Antifade Reagent containing DAPI (Molecular Probes).

### Flow Cytometry

Cultured fibroblasts were washed with PBS and resuspended in PBS containing LipidTOX (1:200). Following incubation for 30 min, cells were subjected to flow cytometry using Accuri C6 (BD Biosciences). Cultured PCLS and/or finely minced mouse lung tissues underwent digestion with 0.5% collagenase (Gibco) in PBS for 45 min at 37 °C while rotating. Cell suspensions were then aspirated through 18, 20, 24G needles and passed through 70- and 40-µm cell strainers (Greiner Bio-One). Cells were pelleted, resuspended in PBS and stained with anti-CD45 (Biolegend, 1:100), CD31 (Biolegend, 1:100) and CD326 (EpCAM, Biolegend, 1:50) antibodies, as well as LipidTOX (1:200). Stained cell suspensions were then analyzed using LSRFortessa (BD Biosciences). Data were analyzed using FlowJo software (FlowJo, LLC).

### Gene expression microarrays

Total RNA Cy5-labeling was carried out using the LIRAK kit (Agilent Technologies) according to manufacturer’s instructions. Per reaction, 200 ng of total RNA was used. Cy-labeled aRNA was hybridized overnight to 8 x 60K 60mer oligonucleotide-spotted microarray slides (Agilent Technologies, design ID 028005). Hybridization and subsequent washing and drying of the slides were performed following the Agilent hybridization protocol. Dried slides were scanned at 2 µm/pixel resolution using the InnoScan is900 (Innopsys). Image analysis was performed using Mapix 6.5.0 software and calculated values for all spots were saved as GenePix results files. Stored data were evaluated using the R software and the limma package from BioConductor. Log mean spot signals were taken for further analysis. Data were quantile-normalized before averaging. Genes were ranked for differential expression using a moderated t-statistic. Pathway analyses were done using gene set tests on the ranks of t-statistics.

### Kinase activity assay

Kinase activity of metformin- or vehicle-treated IPF fibroblasts (n=3 per group) was analyzed by the PamStation (PamGene International BV) that uses a methodology that allows robust analysis of the activity of tyrosine as well as serine/threonine kinases in cells and tissues^23–26^. Hereby, active kinases phosphorylate their distinct peptide substrates presented on a peptide array chip. Phosphorylated peptides are recognized by phospho-specific FITC-labeled antibodies and detection, performed in multiple cycles at different exposure times, is monitored by a CCD camera. Software-based image analysis integrates the signals obtained within the time course of the incubation of the kinase lysate on the chip into one single value for each peptide for each sample (exposure time scaling). Log transformation of processed signals allows easier graphical presentation of the raw data. Thereby, data with significant differences in intensity are visualized on the log-transformed y-axis in a heat map that shows the degree of phosphorylation for each peptide.

For protein isolation including active kinases, IPF fibroblasts were washed with 5 mL ice-cold PBS and scraped from the dishes in 100 µL of M-PER lysis buffer (Thermo Fisher Scientific) containing protease and phosphatase inhibitor cocktails (Pierce). The lysate was incubated for 1 h at 4°C with constant agitation followed by centrifugation at 16,000 g for 15 min at 4°C. The supernatant was immediately flash-frozen in liquid nitrogen and stored at -80°C. Protein concentration was determined using a bicinchoninic acid (BCA) protein assay kit (Thermo Fisher Scientific) according to manufacturer’s instructions.

For tyrosine kinase activity detection, 10 µg of protein lysate was dispensed onto an array of the PamChip PTK (phospho-tyrosine kinase) dissolved in protein kinase buffer and additives (proprietary information) including 1% BSA, 10 mM DTT, 0.6 µL FITC-conjugated antibodies and 400 µM ATP in a final volume of 40 µL (assay master mix). A total of 1 µg of protein lysate was used for serine/threonine kinase activity detection on an array of the PamChip STK (serine/threonine kinase) with protein kinase buffer (proprietary information) supplied with 1% BSA, 0.46 µl primary STK antibody mix and 400 µM ATP (sample mix). After an initial incubation time, secondary FITC-labeled antibodies (0.4 µL) were added. The mixture was dissolved in antibody buffer (proprietary information) and water in a final volume of 30 µL (detection mix).

Upstream kinase prediction on the basis of the different phosphorylation pattern in metformin- and vehicle-treated IPF fibroblasts was conducted using the Bionavigator software v.6.3.67.0 (PamGene International).

### Statistical analyses

Statistical analyses and graph assembly were carried out using GraphPad Prism 6 (GraphPad Prism Software). Outliers were identified using the ROUT method and D’Agostino-Pearson normality test was applied. Student’s t-test (unpaired, two-tailed) was utilized to compare the means of two groups, while multiple pair-wise comparisons were used to compare the means of three or more groups. The number for biological samples (n) for each group is stated in the corresponding figure legend. Differences in means were considered statistically significant if P<0.05.

## Results

### Metformin induces lipid-droplet accumulation in human IPF lung fibroblasts

In order to investigate whether metformin impacts lipogenic versus myogenic fibroblastic phenotypes in the lung, we carried out a dose-finding study. Human lung fibroblasts isolated from 7 IPF patients were starved for 24 h before being treated with 1, 5 or 10 mM metformin for 72 h (Fig. S1). Treatment of cells with 5 mM metformin resulted in significant upregulation of the lipogenic markers adipose differentiation-related protein (a.k.a. perilipin 2, *PLIN2*, 18.3 folds, Fig. S1A) and *PPARg* (6.6 folds, Fig. S1B) in parallel to a significant 9.7-fold downregulation of *COL1A1* (Fig. S1C). Treatment with 10 mM metformin led to comparable results (Fig. S1A-C). Therefore, we decided to use the 5 mM concentration for the rest of the study.

In order to gain further insights into the dynamics of this process, human lung fibroblasts isolated from 5 IPF patients were treated with 5 mM metformin and analyzed after 48, 72 and 96 h (Fig. S2A). Analysis at 48 h showed a significant 3.4-fold downregulation of *COL1A1* expression (Fig. S2D) but without affecting lipogenic marker expression (Fig. S2B, C) or lipid-droplet accumulation (as shown by staining with the neutral lipid dye LipidTOX) (Fig. S2K, L, Q). At 72 h, a significant 9.6-fold downregulation of *COL1A1* (Fig. S2G) was accompanied by significant upregulation of *PLIN2* (33.5 folds, Fig. S2E) and *PPARg* (12.4 folds, Fig. S2F) and a 4.3-fold increase in lipid-droplet accumulation (Fig. S2M, N, R). The effect of metformin was comparable at 96 h (Fig. S2H-J, O, P, S). These data indicate that metformin first leads to *COL1A1* downregulation and then induces lipogenic marker expression in primary human IPF lung fibroblasts. The 72 h time point was chosen for subsequent analyses.

Following the establishment of an optimal dose and time point for metformin treatment, pulmonary fibroblasts isolated from 12 IPF patients were treated with 5 mM metformin and analyzed after 72 h (Fig. 1A). The results showed significant upregulation of *PLIN2* (22.1 folds, Fig. 1B) and *PPARg* (8.3 folds, Fig. 1C), and a robust 8.1-fold downregulation of *COL1A1* (Fig. 1D). Despite the variation in the response of fibroblasts isolated from different patients, the pattern of response was highly significant and clearly consistent among all samples (i.e. induction of lipogenic marker expression and suppression of *COL1A1* expression). As a final readout for lipogenic differentiation, the neutral lipid stain LipidTOX was used and the increase in lipid-droplet accumulation in fibroblasts was demonstrated by fluorescence microscopy (Fig. 1E, F). Flow cytometry-based quantification of LipidTOX^+^ cells displayed a significant 2.1-fold increase in metformin-treated cells compared to vehicle-treated cells (Fig. 1G-I).

**Figure 1.**
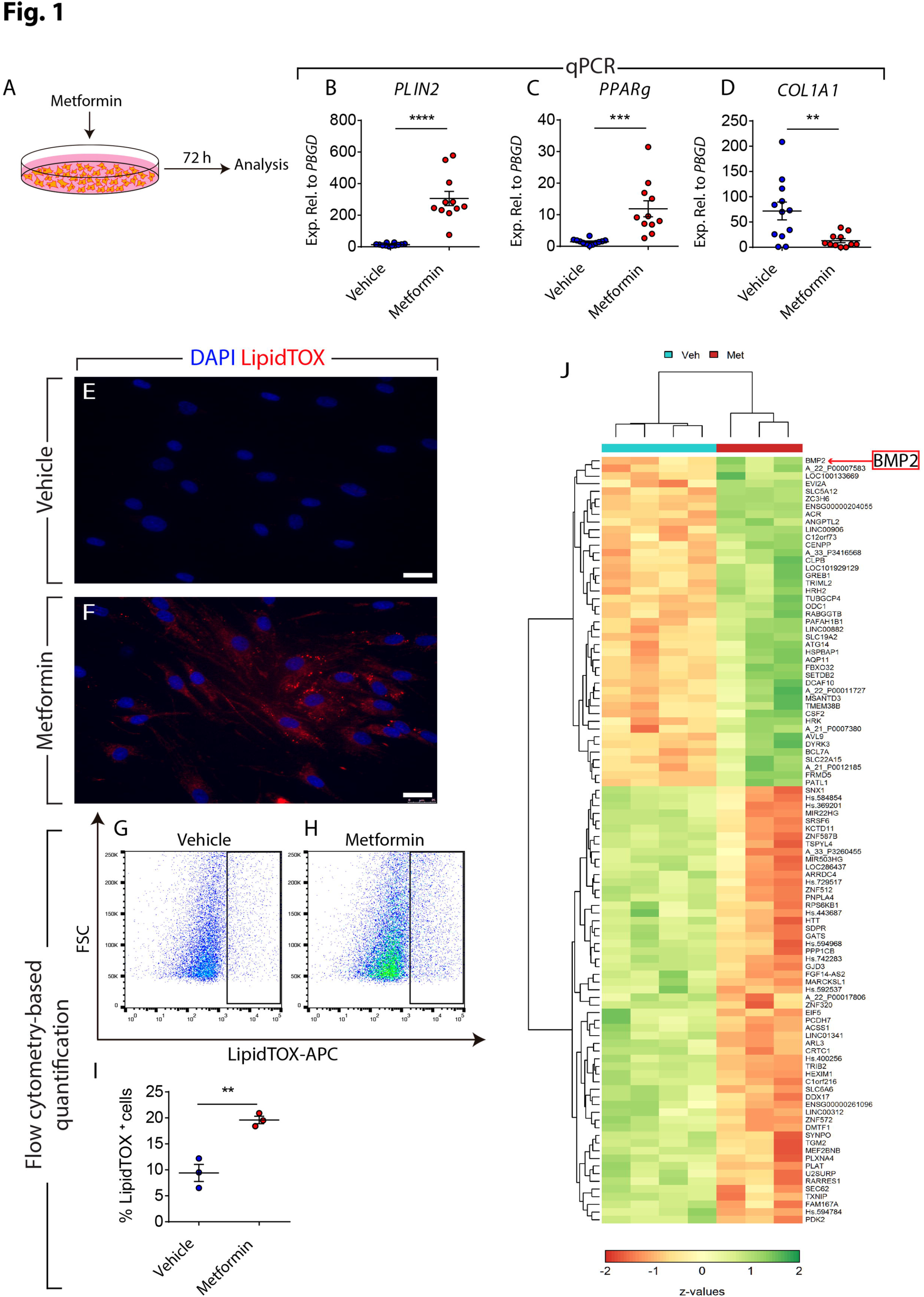
Metformin induces lipogenic marker expression in human IPF lung fibroblasts. **(A)** Schematic representation of the experimental setup. **(B-D)** qPCR analysis for the lipogenic marker genes *PLIN2* and *PPARg*, as well as *COL1A1* in human lung fibroblasts treated with metformin or vehicle. **(E, F)** Staining of lipid droplets in fibroblasts using LipidTOX (red). Nuclei were counterstained with DAPI (blue). **(G-H)** Gating strategy for detecting LipidTOX^+^ cells by flow cytometry. **(I)** Quantification of LipidTOX^+^ cells in response to metformin treatment. **(J)** Heatmap representation of the top 100 differentially expressed genes in fibroblasts following metformin treatment. Scale bars: (E, F) 25 µm. (B-D) Each data point within a given group corresponds to one patient. Vehicle-treated group: n=12, Metformin-treated group: n=11. (I) n=3 per group. ** P<0.01, *** P<0.001, **** P<0.0001.

Next, gene expression microarrays were carried out for 4 vehicle-treated and 3 metformin-treated IPF lung fibroblasts. A heatmap depicting the top 100 regulated genes (based on z-values) are shown in Fig. 1J. Interestingly, KEGG pathway analysis showed that the most significantly downregulated metabolic pathway upon metformin treatment was steroid biosynthesis. Another significantly downregulated pathway was glycosaminoglycan biosynthesis - heparan sulfate / heparin, which has been shown to be enhanced in IPF^27,28^. Other significantly downregulated pathways included N-Glycan biosynthesis, metabolism of xenobiotics by cytochrome 450, drug metabolism – cytochrome P450, retinol metabolism, vitamin B6 metabolism, terpenoid backbone biosynthesis and unsaturated fatty acid biosynthesis.

Interestingly, metformin-treated cells also displayed upregulation of genes involved in sphingolipid metabolism (although less significantly) and downregulation of genes involved in arginine and proline metabolism. This corresponds to previous findings in human IPF lungs, reported to be characterized by reduced sphingolipid metabolism and increased arginine and proline metabolism^14^. The latter pathway is important for collagen synthesis. Other downregulated signaling pathways included TGFβ, ECM-receptor interactions, protein processing in the endoplasmic reticulum, vascular and cardiac smooth muscle contraction, glycolysis / gluconeogenesis and insulin resistance. Upregulated pathways included insulin secretion, regulation of autophagy, lysine degradation and linoleic acid metabolism. Intriguingly, the top upregulated gene in metformin-treated fibroblasts was *BMP2* (red arrow in Fig. 1J).

### Metformin inhibits **TGF**β1-mediated fibrogenesis in vitro

The TGFβ1 signaling pathway is widely regarded as the master regulator of fibrogenesis in vitro and in vivo^29^. In order to investigate whether metformin inhibits the profibrotic effect of TGFβ1 in vitro, primary lung fibroblasts isolated from 10 IPF patients were treated with 2 ng/mL rhTGFβ1 (or vehicle) for 72 h and then treated with 5 mM metformin (or vehicle) for 72 h (Fig. 2A). This experimental setup allows examining the ability of metformin to change the phenotype of myofibroblasts that arise from TGFβ1 treatment. The effect of TGFβ1 treatment was validated after 72 h by qPCR and the results showed significant downregulation of the lipogenic markers *PLIN2* (Fig. 2B) and *PPARg* (Fig. 2C) in parallel to significant upregulation of *COL1A1* (Fig. 2D) as we previously described^8^. We also expected that activation of TGFβ1 signaling would result in disappearance of lipid droplets at the expense of *ACTA2* and *COL1A1* upregulation. To test this hypothesis, cells were stained with LipidTOX and anti-ACTA2 antibodies. Fluorescent microscopy showed increased abundance of ACTA2 filaments (Fig. 2H) and absence of lipid droplets (Fig. 2F) in TGFβ1-treated cells compared to vehicle-treated cells (Fig. 2E, G) after 72 h. Time-lapse microscopy of vehicle- (supplementary movie 1), rhTGFβ1- (supplementary movie 2) or metformin-treated cells (supplementary movie 3) in the presence of LipidTOX was also carried out for 65 h. Metformin led to an increase in lipid-droplet accumulation after 48 h while vehicle or rhTGFβ1 treatment led to cell elongation and disappearance of lipid droplets.

**Figure 2.**
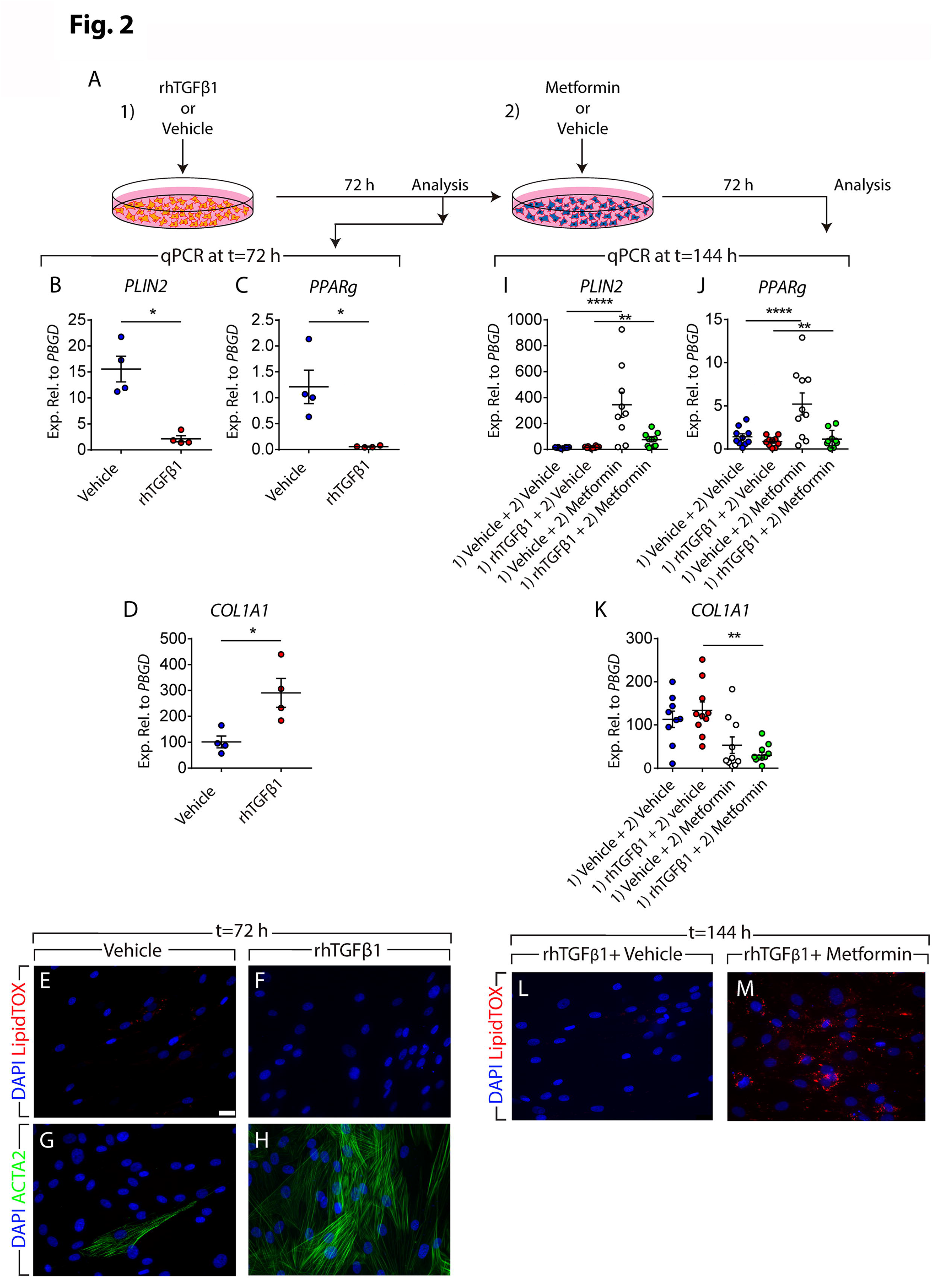
Metformin attenuates TGFβ1-mediated fibrogenesis in vitro. **(A)** Schematic representation of the experimental setup. **(B-D)** qPCR analysis for *PLIN2, PPARg* and *COL1A1* in human lung fibroblasts treated with TGFβ1 or vehicle for 72h. **(E-H)** Staining of TGFβ1- and vehicle-treated cells with LipidTOX (red), anti-ACTA2 antibodies (green) and DAPI (blue). **(I-K)** qPCR analysis for *PLIN2, PPARg* and *COL1A1* in human lung fibroblasts treated with TGFβ1 or vehicle for 72 h, followed by treatment with metformin or vehicle for 72 h. **(L-M)** Staining of TGFβ1-and vehicle-treated cells with LipidTOX (red) and DAPI (blue) at the end of treatment (t=144 h). Scale bars: (E-H) and (L-M) 25 µm. (B-D, I-K) Each data point within a given group corresponds to one patient. (B-D) n=4 per group. (I-K) n=9-10 per group. * P<0.05, **P<0.01, ****P<0.0001.

Next, the response to metformin treatment was analyzed. At the end of day 6 (144 h), qPCR showed significant upregulation of *PLIN2* (4.6 folds, Fig. 2I) and *PPARg* (1.3 folds, Fig. 2J), and a significant 7.1-fold downregulation of *COL1A1* expression (Fig. 2K) in the [TGFβ1 plus metformin] group compared to the [TGFβ1 plus vehicle] group. Interestingly, lipid-droplet accumulation was regained in fibroblasts treated with [TGFβ1 plus metformin] compared to the [TGFβ1 plus vehicle] group (Fig. 2L,M). These results suggest that metformin affects the phenotype of myofibroblasts arising from TGFβ1 treatment by downregulating myogenic markers and inducing lipogenic differentiation.

### Metformin improves lung structure in an ex vivo system

One drawback of using cell-culture systems is that cells growing in vitro are deprived of their microenvironmental cues and their behavior might not resemble that of an in vivo context. We therefore set out to test whether metformin exerts similar beneficial effects in a more complex system that better mimics the in vivo setting. Therefore, we used precision-cut lung slices (PCLS), an ex vivo culture system that has been previously described^30^. This technique allows maintenance of viable, metabolically active lung tissue with preserved structure for five days^30^. Two hundred micrometer-thick PCLS were prepared from fibrotic regions of fresh IPF lung tissues and were cultured in DMEM supplemented with 10% BCS and in the presence or absence of 5 mM metformin for five days (Fig. 3A). Bright-field imaging showed that metformin-treated PCLS displayed a more relaxed structure with open airspaces (Fig. 3E) compared to their vehicle-treated counterparts (Fig. 3D). PCLS were then embedded in paraffin, sectioned into thin slices, and processed for histological analysis. Hematoxylin and eosin staining (Fig. 3H, I), Masson’s trichrome staining (Fig. 3J, K) and COL1A1 immunostaining (Fig. 3L, M) showed improved lung structure and decreased collagen deposition in metformin-treated samples compared to controls. Another set of PCLS was not sectioned but was subjected to whole-mount staining using anti-COL1A1 antibodies and LipidTOX followed by confocal microscopy. Three-dimensional (3D) reconstruction using z-stacks acquired from these samples revealed decreased COL1A1 expression and increased lipid-droplet accumulation (Fig. 3N, O). Flow cytometry-based quantification confirmed the histological observations and showed a significant increase in the abundance of lipid-droplet-containing cells in metformin-treated PCLS (12 ± 0.6%) compared to controls (7.2 ± 1.1%) (Fig. 3P, Q). Finally, total collagen assay showed a significant reduction in collagen content from 26.9 ± 0.9 µg/µL to 20.8 ± 0.9 µg/µL in response to metformin treatment (Fig. 3R).

**Figure 3.**
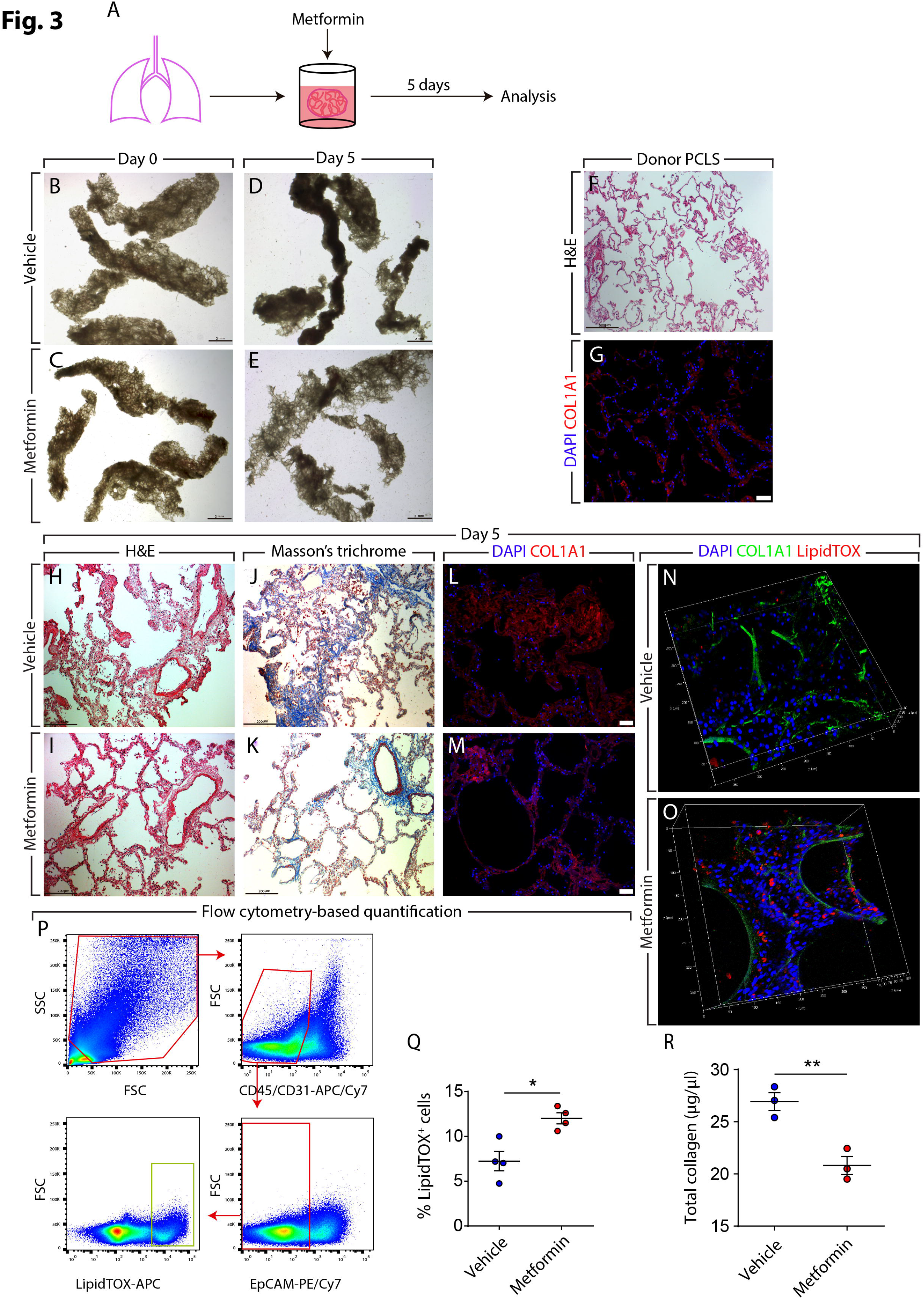
Metformin improves IPF lung structure ex vivo. **(A)** Schematic representation of the experimental setup. **(B-E)** Bright-field imaging of PCLS treated with metformin or vehicle for five days. **(F, G)** Hematoxylin and eosin staining and COL1A1 immunostaining of PCLS prepared from a non-IPF donor lung. **(H-M)** Hematoxylin and eosin staining, Masson’s trichrome staining and COL1A1 immunostaining of PCLS prepared from an IPF lung and treated with metformin or vehicle for five days. **(N, O)** 3D-reconstruction of z-stacks of metformin- and vehicle-treated PCLS stained for COL1A1 (green) and lipid droplets (red). **(P)** Gating strategy for flow cytomety-based quantification of LipidTOX^+^ cells that are negative for hematopoeitic (CD45), endothelial (CD31) and epithelial (EpCAM) cell markers. **(Q)** Quantification of flow cytometry measurements on metformin- and vehicle-treated cells. **(R)** Total collagen assay for metformin- and vehicle-treated cells. Scale bars: (B-E) 2 mm, (F) 500 µm, (G, L, M) 50 µm, (H-K) 200 µm. (Q, R) Each data point within a given group corresponds to one patient. (Q) n=4 per group. ^®^ n=3 per group. * P<0.05, **P<0.01.

### Metformin accelerates resolution of bleomycin-induced pulmonary fibrosis in mice

We next sought to test whether the ability of metformin to alter the myofibroblast fate also applies to the in vivo context of lung fibrosis. In order to address this question, we employed a lineage-tracing approach in the context of bleomycin-induced pulmonary fibrosis (Fig. 4). This model of injury is reversible and animals start to spontaneously recover from lung fibrosis after day 14. Moreover, we previously reported that the *Acta2-Cre-ERT2; tdTomato^flox^* lineage-tracing tool allows genetic labeling of ACTA2^+^ myofibroblasts as they accumulate during the buildup of lung fibrosis when tamoxifen is introduced between days 5 and 14 after bleomycin injury^8^. Therefore, double transgenic animals were treated with bleomycin, fed tamoxifen-containing pellets between days 5 and 14, treated with metformin (or vehicle) starting at day 14 after bleomycin instillation, and were sacrificed at day 28 (Fig. 4A, B). This experimental setup allows labeling myofibroblasts that form during fibrosis formation and tracing their fate during resolution in the presence or absence of metformin. Already upon gross optical inspection, hematoxylin and eosin as well as masson’s trichrome stained lungs explanted at day 28 showed enhanced recovery from fibrosis upon metformin treatment (Fig. 4C-F). Correspondingly, quantification of lung fibrosis showed a significant decrease in the extent of fibrosis from 12 ± 1.5% in the vehicle-treated group to 8 ± 0.6% in the metformin-treated group (Fig. 4G). Immunofluorescent staining allowed visualization of lineage-traced myofibroblast-derived cells (tdTomato^+^) and also confirmed the decrease in COL1A1 deposition upon metformin treatment (Fig. 4H, I). Interestingly, LipidTOX staining carried out on frozen sections revealed the presence of lipid droplets in tdTomato^+^ cells at day 28 (Fig. 4J, K). In order to quantify the impact of metformin on altering the myofibroblast fate, flow cytometry was carried out. The results showed similar proportions of tdTomato^+^ cells in both groups (Fig. 4Q), while the proportion of LipidTOX^+^ mesenchymal cells increased from 13.5 ± 2.5% to 18.8 ± 1.8% upon metformin treatment (Fig. 4S). More importantly, the proportion of myofibroblast descendants (tdTomato^+^) that also contained lipid droplets increased from 6.8 ± 0.1% in the control group to 12.2 ± 0.5% in the metformin-treated group (Fig. 4R). Collectively, these data clearly demonstrate that metformin accelerates fibrosis resolution and this is accompanied by the induction of myogenic-to-lipogenic conversion in lung fibroblasts in vivo.

**Figure 4.**
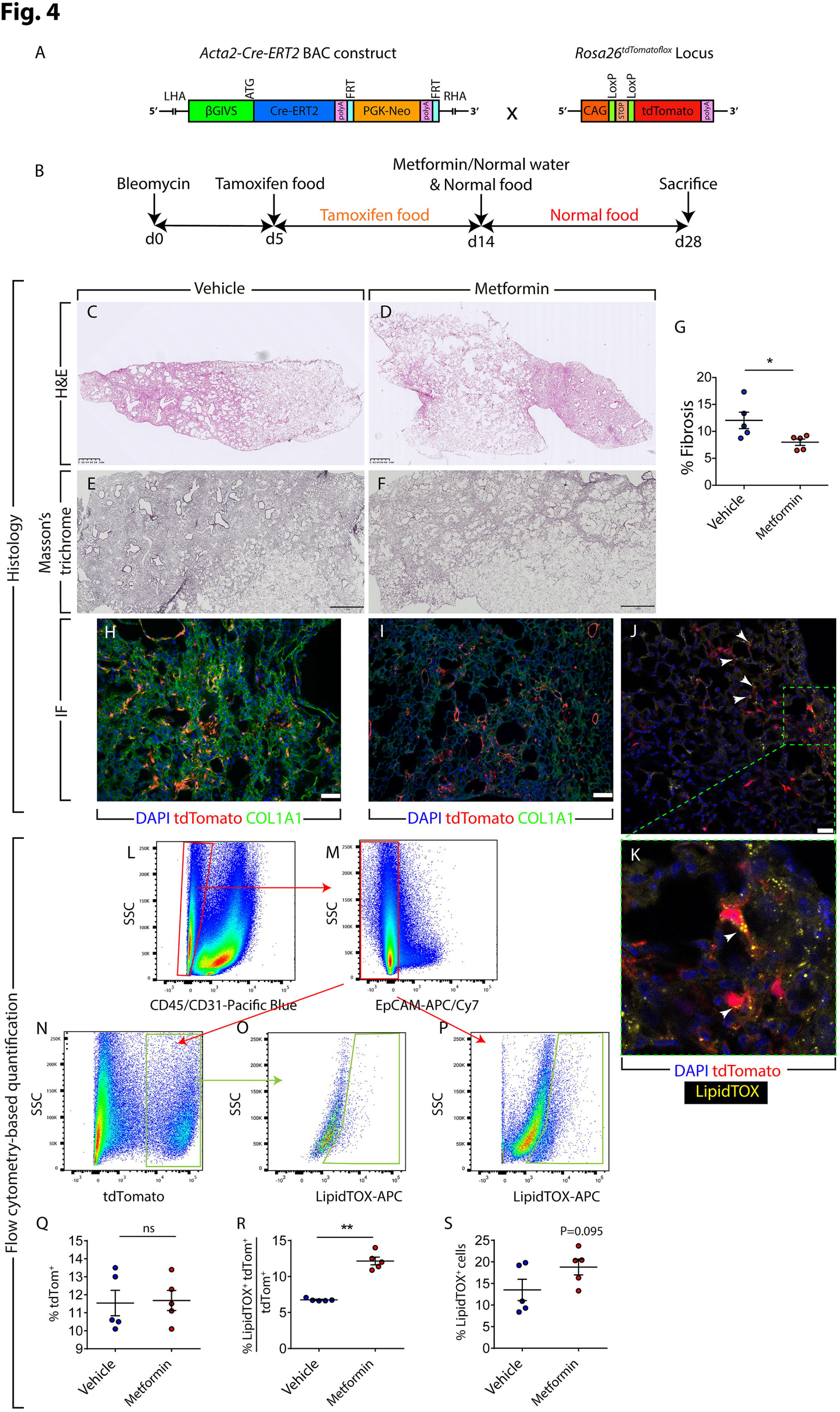
Metformin accelerates fibrosis resolution in the bleomycin model in mice. **(A)** Schematic representation of the *Acta2-Cre-ERT2* and *tdTomato^flox^* construct. **(B)** Schematic representation of the timeline of the experiment. Bleomycin was administered intratracheally at day 0. Between days 5 and 14, mice were fed tamoxifen-containing pellets and starting at day 14, metformin (1.5 mg/mL) or vehicle was administered through drinking water. Mice were sacrificed at day 28. **(C-F)** Hematoxylin and eosin and Masson’s trichrome staining of metformin- and vehicle-treated lungs. **(G)** Quantification of fibrosis in metformin- and vehicle-treated lungs. **(H, I)** Immunofluorescence for COL1A1 (green). Endogenous tdTomato signal (red) and DAPI (blue) are also shown. **(J)** LipidTOX staining (green) and tdTomato^+^ cells (red) are shown. The box in (J) is magnified in **(K).** Arrowheads indicate LipidTOX^+^ tdTomato^+^ cells. **(L-S)** Gating strategy (to detect CD45^-^ CD31^-^ EpCAM^-^ tdTomato^+^ and/or LipidTOX^+^ cells) and quantification of various cell populations based on tdTomato and LipidTOX detection. Scale bars: (C-F) 1 mm, (H, I) 50 µm, (J) 25 µm. (G, Q-S) Each data point within a given group corresponds to one animal. n=5 per group. * P<0.05, **P<0.01. IF: Immunofluorescence, ns: Not significant.

### The mechanism of action of metformin in human lung fibroblasts is only partially dependent on AMPK signaling

Activation of the AMPK signaling cascade is one of the major molecular mechanisms that have been proposed for metformin. In order to investigate whether metformin-mediated lipogenic differentiation is AMPK-dependent, gain and loss-of-function approaches were carried out. In a first set of experiments, primary human lung fibroblasts isolated from 8 IPF patients were cultured and treated with GSK621 (a selective activator of AMPK signaling pathway) for 72 h (Fig. 5A). The results showed a trend for mild downregulation of the lipogenic markers *PLIN2* (Fig. 5B) and *PPARg* (Fig. 5C) in parallel to a robust, significant downregulation of *COL1A1* (Fig. 5D). Accordingly, GSK621 treatment failed to promote lipid-droplet accumulation in these cells (Fig. 5L-N).

**Figure 5.**
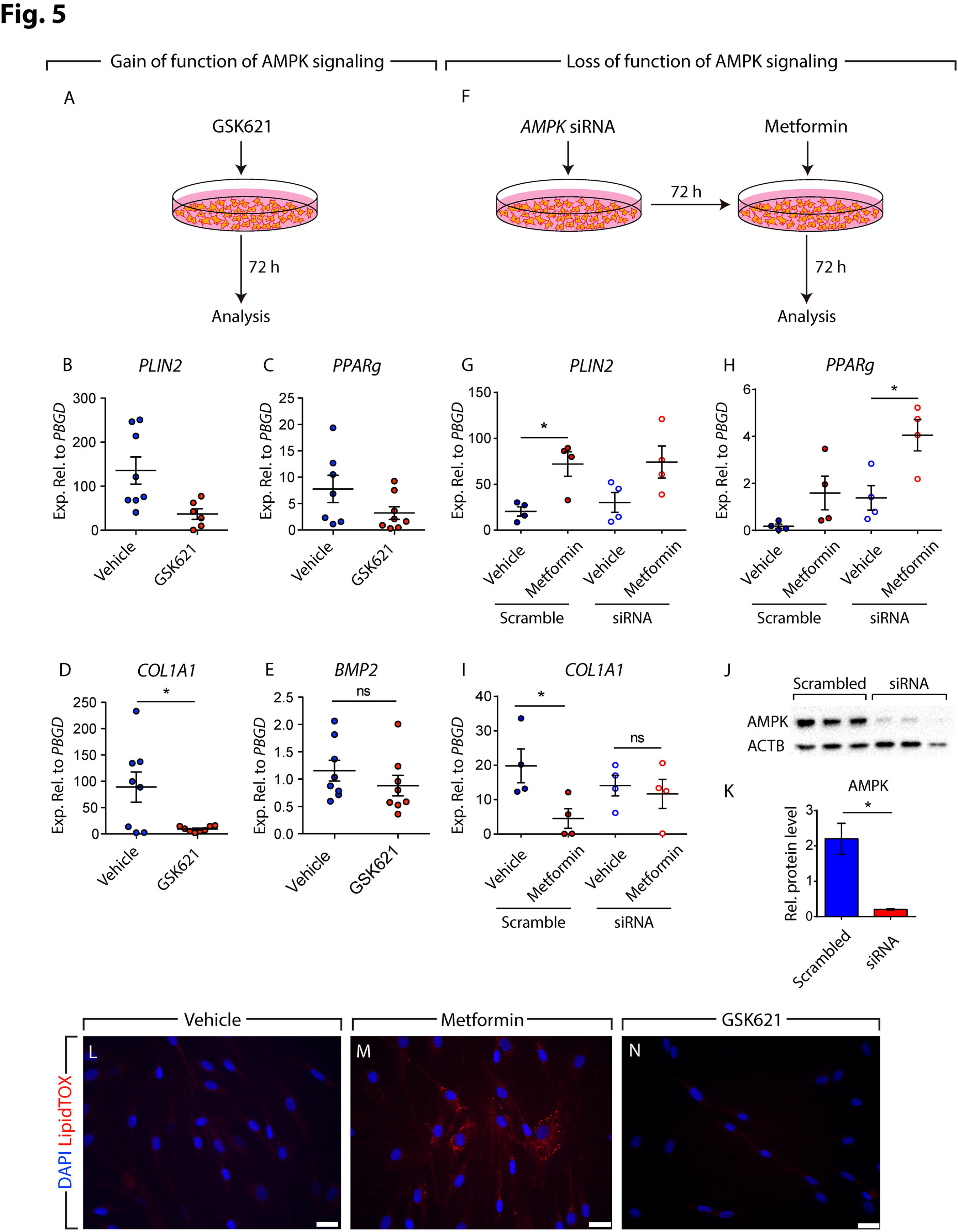
Mode of action of metformin is partially independent of AMPK signaling. **(A)** Schematic representation of the gain-of-function experimental setup for AMPK signaling. **(B-E)** qPCR analysis of *PLIN2, PPARg*, *COL1A1* and *BMP2* in IPF fibroblasts treated with AMPK agonist GSK621 or vehicle. **(F)** Schematic representation of the loss-of-function experimental setup for AMPK signaling. **(G-I)** qPCR analysis of *PLIN2, PPARg* and *COL1A1* in IPF fibroblasts treated with *AMPK* siRNA or scramble siRNA. The decrease of AMPK protein levels at the time of analysis is shown in **(J, K). (L-N)** Staining of GSK621- and vehicle-treated cells with LipidTOX (red) and DAPI (blue). Metformin-treated cells were used as a positive control for lipid-droplet accumulation (M). Scale bars: (L-N) 25 µm. (B-E, G-I, K) Each data point corresponds to one patient. (B-E) Vehicle-treated group: n=7-8, GSK621-treated group: n=6-8. (G-I) n=4 per group. (K) n=3 per group. * P<0.05. ns: Not significant.

In another set of experiments, primary human lung fibroblasts isolated from 5 IPF patients were transfected with siRNAs targeting *AMPK*. After 72 h, cells were treated with metformin and were analyzed after 72 h (Fig. 5F). Western blotting revealed a 90% knockdown of *AMPK* in these cells compared to scramble-transfected cells at the time of analysis (Fig. 5J, K). Interestingly, metformin treatment led to upregulation of *PLIN2* and *PPARg* regardless of *AMPK* knockdown (Fig. 5G, H). Metformin-mediated *COL1A1* downregulation, however, was attenuated upon *AMPK* knockdown (Fig. 5I). Altogether, these data demonstrate that the mechanism of action of metformin in promoting lipogenic differentiation in human lung fibroblasts is largely independent of AMPK signaling while inhibition of COL1A1 production is mainly AMPK-dependent.

### Metformin induces the expression of lipogenic markers via induction of *BMP2* expression and phosphorylation of **PPAR**γ

The next step was to better define the molecular mechanism by which metformin induces lipogenic differentiation in human lung fibroblasts. Our gene expression microarrays showed that *BMP2* was the highest upregulated gene in fibroblasts upon metformin treatment (Fig. 1J). BMP2 is known to inhibit smooth muscle cell growth in a mechanism involving PPARγ activation^31^. Calvier et al. also reported that BMP2 inhibits TGFβ1 signaling via PPARγ in vascular smooth muscle cells in the lung^32^. Therefore, we treated 11 human IPF lung fibroblasts with rhBMP2 and gene expression was analyzed after 72 h (Fig. 6A). Intriguingly, rhBMP2 treatment resulted in significant upregulation of *PLIN2* (1.9 folds, Fig. 6B) and *PPARg* (2.5 folds, Fig. 6C), while *COL1A1* expression levels remained unchanged (Fig. 6D). LipidTOX staining confirmed the increase in lipid-droplet accumulation in these cells (Fig. 6E, F). Thus, BMP2 is a positive regulator of lipogenic differentiation in human IPF lung fibroblasts.

**Figure 6.**
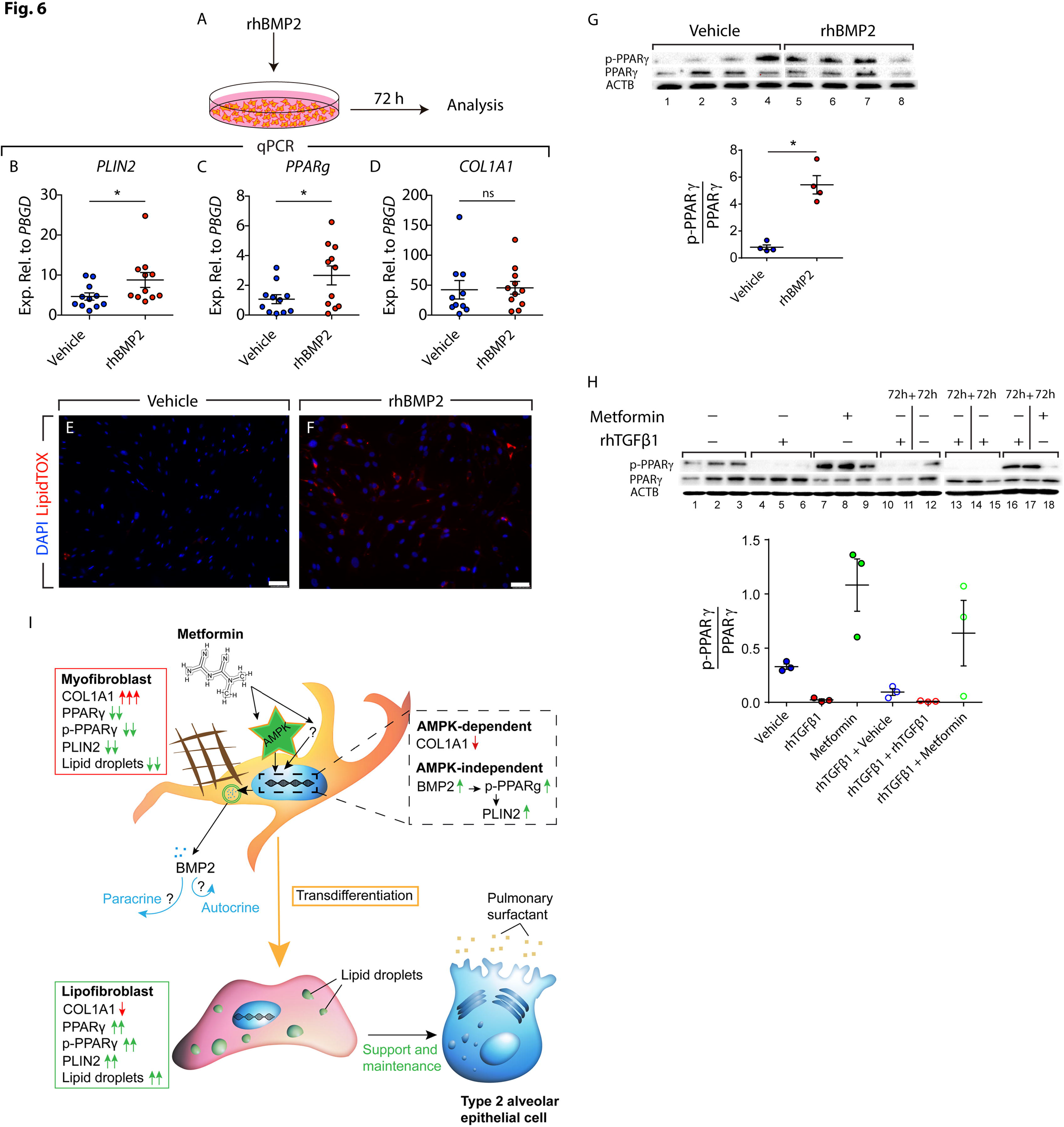
rhBMP2 and metformin induce PPARγ phosphorylation and lipogenic differentiation in human IPF lung fibroblasts. **(A)** Schematic representation of the experimental setup. **(B-D)** qPCR analysis of *PLIN2, PPARg* and *COL1A1* in IPF fibroblasts treated with rhBMP2 or vehicle. **(E, F)** Staining of rhBMP2- and vehicle-treated cells with LipidTOX (red) and DAPI (blue). **(G)** Western blot showing the induction of PPARγ phosphorylation in response to rhBMP2 treatment. Lanes 1-4 and lanes 5-8 were run in parallel on different gels under the same conditions. Quantification of the immunoblot is shown in the lower panel. **(H)** Western blot showing the opposing effects of metformin and TGFβ1 on PPARγ phosphorylation, and the ability of metformin to partially restore PPARγ phosphorylation in TGFβ1-treated cells. Lanes 1-12 and lanes 13-18 were run in parallel on different gels under the same conditions. Quantification of the immunoblot is shown in the lower panel. **(I)** Model for the antifibrotic mechanism of action of metformin in human lung fibrosis. Metformin activates AMPK signaling in myofibroblasts, leading to suppression of collagen production, and induces lipogenic differentiation via an AMPK-independent mechanism involving BMP2 release and PPARγ activation. Arising lipofibroblasts are known to support type 2 alveolar epithelial stem cells in the lung. Scale bars: (E-F) 50 µm. (B-D, G, H) Each data point corresponds to one patient. (B-D) n=10-11 per group. (G) n=4 per group. (H) n=3 per group. * P<0.05, ns: Not significant.

The activation of PPARγ signaling pathway can also be induced post-translationally via PPARγ phosphorylation^33^. Therefore, we set out to determine whether metformin and BMP2 induce lipogenic differentiation in human lung fibroblasts by inducing PPARγ phosphorylation. Firstly, cells were treated with rhBMP2 (or vehicle) and protein lysates were collected after 72 h. PPARγ phosphorylation was induced 7.6 folds in rhBMP2-treated fibroblasts (Fig. 6G). In another set of experiments, treatment of cells with TGFβ1 led to a 24.4-fold reduction in p-PPARγ levels (Fig. 6H, lanes 4-6) while treatment with metformin led to a 3.3-fold increase in p-PPARγ levels (Fig. 6H, lanes 7-9) compared to vehicle-treated cells (Fig. 6H, lanes 1-3). In order to investigate whether metformin counteracts the profibrotic effects of TGFβ1 in fibroblasts by modulating the phosphorylation status of PPARγ, cells were first treated with rhTGFβ1 for 72 h and then followed by metformin treatment for another 72 h. The results showed that transient or continuous treatment with rhTGFβ1 maintained the phosphorylation status of PPARγ at minimal levels (Fig. 6H, lanes 10-15), while treatment with metformin after rhTGFβ1 treatment rescued p-PPARγ levels significantly (Fig. 6H, lanes 16-18). Of note, qPCR analysis revealed that activation of AMPK signaling via GSK621 treatment did not induce *BMP2* expression in IPF fibroblasts (Fig. 5E), indicating that the metformin-BMP2-p-PPARγ axis is independent of AMPK signaling.

### Pirfenidone and nintedanib do not induce lipogenic differentiation in human lung fibroblasts

Currently, there are two FDA-approved drugs for treating IPF; pirfenidone, which is believed to act as a TGFβ1 inhibitor, and nintedanib, which is a multi-receptor tyrosine kinase inhibitor. Therefore, we set out to determine whether these two drugs alter lung fibroblast fate in a similar fashion to metformin. Firstly, primary human lung fibroblasts were treated with either nintedanib (1 µM) or pirfenidone (0.3 mg/ml) for 72 h (Fig. S3A). Quantitative real-time PCR analysis showed that neither nintedanib nor pirfenidone enhanced the expression of the lipogenic markers *PLIN2 and PPARg* (Fig. S3B, C, E, F). Nintedanib and pirfenidone did, however, lead to a 1.9- and 2.3-fold downregulation of *COL1A1*, respectively (Fig. S3D, G). In line with these data, LipidTOX staining did not reveal any change in lipid-droplet accumulation following nintedanib or pirfenidone treatment (Fig. S3H-J). Finally, whole-mount staining followed by confocal imaging and 3D reconstruction of IPF-derived PCLS (cultured for five days in the presence or absence of nintedanib or pirfenidone) revealed a slight decrease in COL1A1 deposition without an evident change in the abundance of LipidTOX^+^ cells (Fig. S3K-O).

## Discussion

Despite recent advances in our understanding of IPF pathology, there is still no curative treatment for IPF, and the currently available antifibrotic treatment modalities retard, but do not completely stop, the progression of the disease. On the other hand, there is emerging literature about the association between metabolic disorders and IPF incidence. We have recently demonstrated an interconversion between lipogenic and myogenic fibroblastic phenotypes in lung fibrosis, a process that is governed by TGFβ1 and PPARγ signaling pathways^8^. Our group and others had already shown that PPARγ agonist rosiglitazone, which is an antidiabetic agent, counteracts TGFβ1-mediated fibrogenesis in vitro and in vivo^8–10^. In this study, we demonstrate that the first-line antidiabetic drug, metformin, inhibits collagen production in primary human lung fibroblasts and in ex vivo cultured human IPF PCLS and strongly enhances myo- to lipofibroblast transdifferentiation linked with phenotypic recovery from fibrosis. Although the PCLS culture system does not fully recapitulate the in vivo situation, it adds cellular, molecular and matrix complexity compared to standard cell culture techniques and – in our opinion – offers a valuable preclinical analytical tool. Accordingly, therapeutic application of metformin in bleomycin-injured mice resulted in accelerated resolution of fibrosis by altering the fate of fibrosis-associated myofibroblasts and inducing their lipogenic differentiation. Notably, detailed pathway analysis showed that the reduction of collagen synthesis was largely AMPK-dependent, whereas the transdifferentiation of myo- to lipofibroblasts occurred in a BMP2-PPARγ-dependent fashion and was largely AMPK independent.

PPARγ is the master regulator of adipogenesis and it is expressed in various cell types in the human body. Its functions include – in addition to differentiation and maintenance of adipocytes^34^ – regulation of inflammatory responses in macrophages^35^, regulation of osteogenesis^36^ and cell metabolism^37^. Therefore, its role in homeostasis and disease is strictly context-dependent. Interestingly, it has been reported that metformin inhibits the upregulation of *PPARg* and *PLIN2* in mouse hepatocytes in response to fructose diet^38^. Metformin, however, does not alter *PPARg* and *PLIN2* expression levels in animals that receive normal diet^38^. Moreover, it has been shown that metformin inhibits adipogenesis in mesenchymal stem cells^39^ and embryonic mouse preadipocyte cell line 3T3-L1^40^. On the other hand, our data clearly demonstrate that metformin significantly upregulates *PPARg* and *PLIN2* in human lung fibroblasts, induces lipid-droplet acquisition, inhibits COL1A1 deposition and regulates metabolic pathways. Therefore, the biological and physiological outcome of metformin treatment is clearly context and cell type-specific. In our in vivo experiments, metformin was introduced through drinking water and the effects on lung repair were evident based on fibrosis assessment and myofibroblast fate switching. No adverse, systemic side effects of such treatment were observed. Although metformin is safe and well tolerated in humans, it confers the risk of hypoglycemia in case of long-term use, although this risk is still lower than that associated with other antidiabetic agents^41^.

Recently, two independent groups have shown that metformin inhibits the profibrotic effect of TGFβ1 in lung fibroblasts via AMKP activation^20,42^. Our data are in line with these reports, but additionally show that activation of PPARγ signaling pathway is a further important mechanism by which metformin attenuates TGFβ1 signaling cascade. Most importantly, we discovered a central role for BMP2-PPARγ-induced myofibroblast-to-lipofibroblast transdifferentiation in the process of fibrosis resolution in response to metformin. Our gain-of-function experiments showed that while AMPK activation significantly downregulates *COL1A1*, it does not induce lipogenic marker expression or lipid-droplet accumulation, which indicates that activation of the AMPK pathway alone is not enough to trigger the transdifferentiation of myo- into lipofibroblasts. Correspondingly, knockdown of *AMPK* did not abolish metformin-mediated induction of lipogenic markers but attenuated the suppression of *COL1A1* expression. These data suggest that alternative signaling mechanisms also contribute to the antifibrotic effects of metformin. Our data strongly suggest that *BMP2* upregulation and PPARγ phosphorylation are centrally involved in these mechanisms. Interestingly, although treatment of primary human IPF lung fibroblasts with rhBMP2 was sufficient to phosphorylate PPARγ and induce lipogenic marker expression, rhBMP2 treatment did not result in *COL1A1* downregulation. Moreover, AMPK activation did not result in *BMP2* upregulation and did not induce major myoto lipofibroblast transdifferentiation. Therefore, we propose a model in which metformin firstly activates AMPK signaling, which downregulates *COL1A1*, and secondly activates an alternative pathway involving *BMP2* upregulation and PPARγ phosphorylation, which induces lipogenic differentiation (Fig. 6I).

Additionally, protein lysates from metformin-treated human IPF fibroblasts were subjected to a protein kinase activity assay (PamStation) in order to identify differentially regulated serine/threonine kinases and tyrosine kinases. Kinases with significantly reduced activities included cyclin-dependent kinase (CDK) family members, MAPK11/14 and ERK1/2 (Fig. S4A). Given the relevance of ERK1/2 in lung fibrosis^43^, we validated the effect of metformin in inhibiting the activity of these kinases by Western blotting and the results showed robust inhibition of p-ERK1/2 levels in IPF lung fibroblasts in response to metformin treatment (Fig. S4B). Treatment of human IPF lung fibroblasts with Selumetinib (a potent and selective inhibitor of MEK that is directly upstream of ERK) did not yield significant changes in the expression levels of lipogenic or myogenic marker genes (Fig. S4D-F). Therefore, we conclude that the ERK pathway is not involved in metformin-mediated myofibroblast-to-lipofibroblast transdifferentiation.

Myofibroblast-to-lipofibroblast trans or redifferentiation is a central hitherto largely underappreciated route for resolution of lung fibrosis. Side-by-side comparison of metformin with pirfenidone and nintedanib showed that the latter agents did not converge on this resolution route. It is worth mentioning that cells and tissues used for pirfenidone, nintedanib or metformin treatment were derived from the same patients. Metformin, like rosiglitazone and maybe other antidiabetic medications, may add a unique antifibrotic profile by inducing transdifferentiation of myo- to lipofibroblasts. Another likely possibility is that since IPF lung tissues are derived from end-stage patients that underwent lung transplantation, it might be that these samples had developed resistance to pirfenidone and nintedanib. Nevertheless, the robust response of these samples to metformin treatment in terms of *COL1A1* downregulation and lipogenic differentiation highlights the therapeutic potential of metformin in IPF.

Many reports have described the involvement of metformin in manipulating various metabolic pathways^44–46^. In fact, metformin is known to disrupt mitochondrial complex I, thus inhibiting cellular respiration. Recently, Zhao et al. published an extensive analysis of metabolic alterations in IPF lungs compared to donors^14^. They concluded that several types of long- and medium-chain fatty acids are enriched in IPF lungs^14^. Sphingolipid metabolism was found to be suppressed while arginine metabolism was found to be enhanced in the IPF lungs^14^. Our KEGG analysis on metformin-treated human lung fibroblasts suggests that metformin might be able to correct such metabolic dysregulations. Better understanding of the metabolic switch in lung fibroblasts in response to metformin and the mechanisms leading to accumulation of lipid droplets warrants further research. Last but not least, the therapeutic role of metformin described in this study might also apply to other fibrotic diseases such as liver fibrosis and scleroderma, characterized by an imbalance between myofibroblasts and adipocyte-like cells (hepatic stellate cells)^47^ and subcutaneous adipocytes^48,49^, respectively.

In a recent report, post-hoc analysis was performed on IPF patients derived from the placebo arms of three phase 3, double-blind, controlled trials of pirfenidone^50^. In that study, 71 metformin users did not present improvements in clinical outcomes compared to 553 non-metformin users^51^. As pointed out by Tzouvelekis and colleagues, these data cannot be generalized into the global IPF population due to many caveats in the study design including the post-hoc nature of the study, the low number of metformin users, lack of stringent criteria for diagnosis and assessment of diabetes control, inability to link metformin mechanisms to IPF pathogenesis or to delineate drug-drug interactions^52^. Given its low cost and the fact that it is well tolerated in humans, it will be useful to test the curative effect of metformin, either alone or in combination with other antifibrotic agents, in non-diabetic IPF patients. A key aspect will be to identify biomarkers that predict drug responsiveness in the heterogeneous population of IPF patients. Given the high financial burden of developing novel drugs, drug repositioning might help accelerate the process of discovering a cure for IPF patients.

To sum up, our data demonstrate a clear antifibrotic role for metformin in the lung. Due to its ability to alter metabolic pathways, to inhibit TGFβ1 signaling, to suppress collagen formation and to promote transdifferentiation of myofibroblasts into lipofibroblasts, metformin should be considered as a novel therapeutic option for IPF patients.

## Acknowledgements

E.E.A. was funded by a start-up grant from the Excellence Cluster Cardio-Pulmonary System (ECCPS). E.E.A. also acknowledges the support of the University Hospital Giessen and Marburg (UKGM) and the German Center for Lung Research (DZL). S.B. was supported by grants from the Deutsche Forschungsgemeinschaft (DFG; BE4443/1-1, BE4443/4-1, BE4443/6-1, KFO309 P7 and SFB1213-projects A02 and A04), Landes-Offensive zur Entwicklung Wissenschaftlich-Ökonomischer Exzellenz (LOEWE), UKGM, Universities of Giessen and Marburg Lung Center (UGMLC), DZL, and COST (BM1201). J.S.Z was funded through a start-up package from Wenzhou Medical University and the National Natural Science Foundation of China (grant number 81472601). S.H. was supported by the UKGM (FOKOOPV), the DZL and grants from the DFG (KFO309 P2/8; SFB1021 C05, SFB TR84 B9).

